# Role of Long Chain Acyl-CoA Synthetases in MASH-driven Hepatocellular Carcinoma and Ferroptosis

**DOI:** 10.1101/2025.03.13.642692

**Authors:** Peyton Classon, Alexander Wixom, Natalia Calixto Mancipe, Rondell P. Graham, Nguyen Tran, Timucin Taner, Davide Povero

## Abstract

Metabolic-associated steatohepatitis-driven hepatocellular carcinoma (MASH-HCC) incidence is rapidly rising worldwide. Lipid metabolic reprogramming is a hallmark of solid tumors to satisfy cancer high metabolic demand. However, it may confer sensitivity to ferroptosis, a cell death mode driven by iron-dependent lipid peroxidation. In this report, we describe the lipid metabolic landscape in MASH-HCC and characterize long chain acyl-CoA synthetases (ACSLs), a family of enzymes involved in synthesis of cellular lipids. Bulk RNA-sequencing, single-cell RNA-sequencing, spatial transcriptomics and immunohistochemistry analyses of human MASH-HCC were integrated to identify differentially expressed lipid metabolism genes. Ferroptosis *in vitro* was assessed in human HCC cell lines. A characterization of ACSLs was also conducted at the single-cell level in a diet-induced experimental murine model of MASH-HCC. Our analysis revealed that in human MASH-HCC, ACSLs exhibit a heterogeneous expression, with ACSL4 notably enriched in tumor tissues, contrasting with ACSL5 upregulation in non-cancerous MASH. We identified a unique lipid metabolic gene signature of MASH-HCC, which included genes associated with ferroptosis vulnerability. *In vitro*, ACSL4 upregulation was associated with increased ferroptosis sensitivity in human HCC cell lines. Lastly, single-cell RNA-sequencing revealed elevated ACSL4 expression in immune cells in a murine MASH-HCC model, suggesting a role of ACSL4 in shaping the tumor immune microenvironment. Overall, this report offers new insights into lipid metabolic landscape and ferroptosis sensitivity for novel MASH-HCC treatments.

**GRAPHICAL ABSTRACT:** 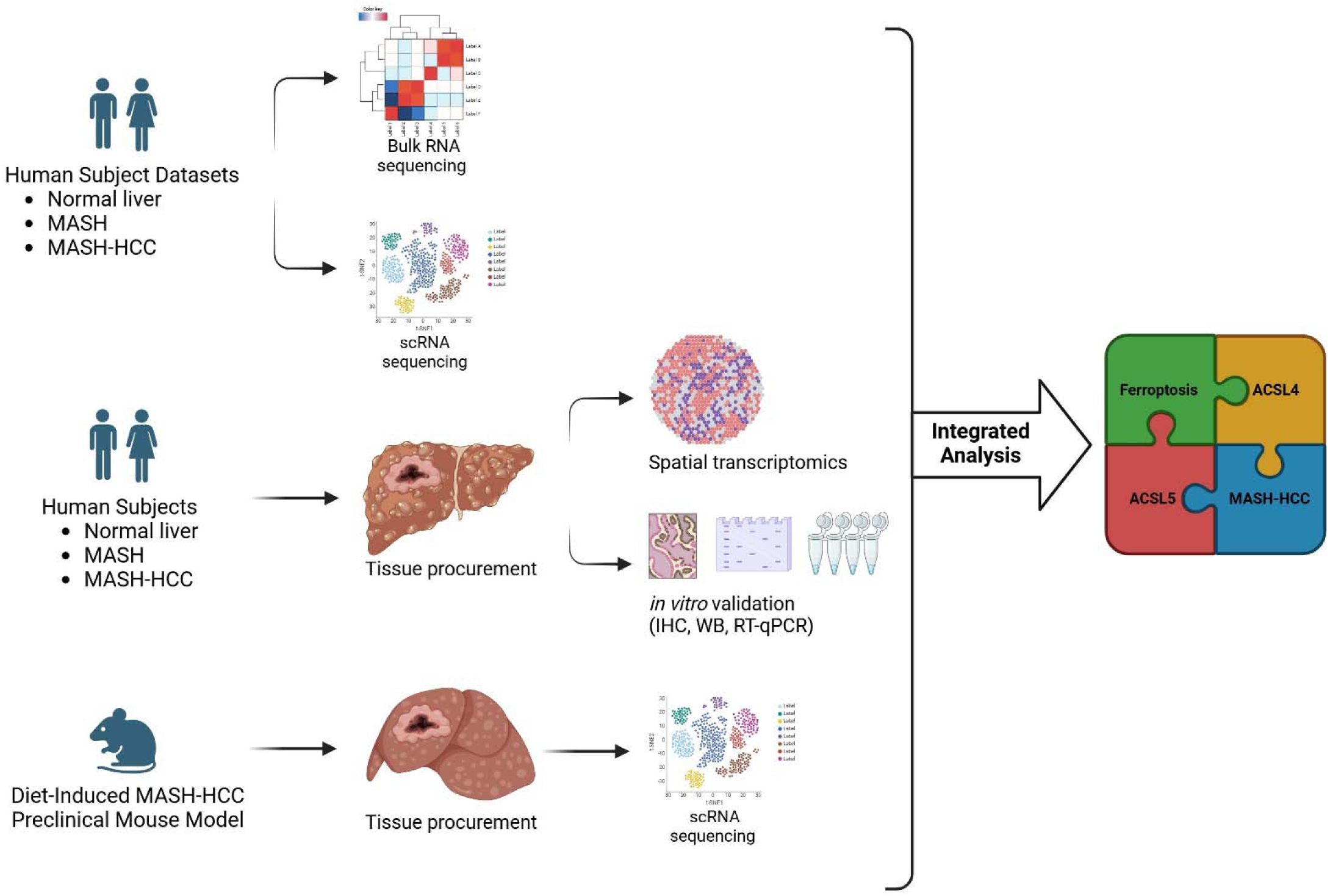

## INTRODUCTION

Metabolic-dysfunction associated steatotic liver disease (MASLD) is one of the most common forms of chronic liver disease, associated with obesity, type II diabetes and metabolic syndrome^1^. MASLD encompasses two sub-types, metabolic-dysfunction associated steatotic liver (MASL), characterized by excessive liver steatosis, and metabolic dysfunction-associated steatohepatitis (MASH), which features steatosis, inflammation, fibrosis and hepatocellular injury^2,3^. When not resolved, MASH can lead to liver cirrhosis and hepatocellular carcinoma (HCC)^4^. MASH-driven HCC (MASH-HCC) is currently the fastest-growing form of HCC^5^.

As a metabolic disease, MASH and MASH-HCC develop in a lipid-rich environment, where lipid metabolic reprogramming occurs to ensure tumor survival, growth and satisfy the tumor metabolic demand^6,7^. As a consequence, expression of genes involved in lipid metabolism differs between the tumor and adjacent non-cancerous areas. Identifying gene expression differences may uncover tumor metabolic vulnerabilities that ensure cancer survival and rapid growth.

Two key strategies utilized by tumors to survive and grow, especially in metabolically stressed conditions, are cell death evasion and metabolic reprogramming. Through metabolic alterations cancer cells obtain the required energy supply or carbon sources for membrane biogenesis to ensure a rapid cell proliferation. However, while advantageous for tumor progression, MASH-HCC lipid metabolic reprogramming may establish a vulnerability to ferroptosis, a novel metabolic, iron-dependent and caspase-independent mode of cell death driven by phospholipid peroxidation.

Three main hallmarks characterize ferroptosis: i) oxidation of polyunsaturated fatty acid-containing membrane phospholipids (PUFA-PL); ii) accumulation of redox-active metal iron; iii) loss of lipid hydroperoxide repair capacity by antioxidant enzymes, such as glutathione peroxidase 4 (GPX4) and ferroptosis suppressor protein 1 (FSP1)^8–10^. Considering that the main driver of ferroptosis is peroxidation of PUFA-rich phospholipids, membrane phospholipid composition and metabolic enzymes involved in phospholipid synthesis and remodeling, dictate the sensitivity to ferroptosis. For example, phospholipids rich in saturated or monosaturated fatty acids are more resistant to lipid peroxidation and ferroptosis, as compared to PUFA-rich phospholipids, the principal targets of peroxidation. A comprehensive and integrated transcriptomic profiling of lipid metabolism in MASH-HCC is currently missing and may not only uncover key insights on metabolic vulnerabilities driving cancer susceptibility to cell death, such as ferroptosis, but also identify novel biomarkers of MASH-HCC.

Based on this, the goal of our report was to characterize and map the lipid metabolic transcriptomics landscape of human and murine MASH-HCC, including non-cancerous MASH and normal liver, utilizing an integrated approach combining bulk and single-cell RNA-sequencing, spatial transcriptomics and targeted *in vitro* and histopathological assays.

Our integrated analysis identified a family of enzymes called Acyl-CoA synthetase long chain (ACSLs). ACSLs include six members that are involved in phospholipid synthesis and fatty acid degradation. Each ACSL activates a different class of long-chain fatty acids and have different subcellular localization. Our findings indicate that: i) ACSLs exhibit a differential and heterogenous expression in human MASH-HCC; ii) ACSL4 is enriched in human MASH-HCC as compared to non-cancerous MASH tissue, where ACSL5 is upregulated; iii) ACSL4 upregulation is associated with increased ferroptosis sensitivity in human HCC cell lines *in vitro*; iv) ACSL4 is upregulated in most immune cells, particularly in natural killer cells (NK) and natural killer T cells (NKT) in an experimental murine model of MASH-HCC, supporting its potential role in shaping the tumor immune microenvironment in MASH-HCC.

In summary, our landscape report on MASH-HCC provides novel insights on lipid metabolism and on the specific molecular and spatial transcriptomics profile of ACSLs, a family of enzymes involved in phospholipid biosynthesis and driving vulnerability to ferroptosis.

## METHODS

### Human Samples

Details on human samples collection and the study population were previously reported^11^. Briefly, MASH-driven HCC tumor tissue and adjacent MASH noncancerous tissues were collected from male and female subjects aged 18 years or older at Mayo Clinic, with biopsy-confirmed MASH-HCC. Tissues were obtained during surgical resection procedures conducted at Mayo Clinic. Individuals with alcohol-associated liver disease, viral hepatitis, or primary sclerosing cholangitis were excluded from the study. Normal liver tissues were procured from subjects undergoing abdominal surgery for reasons unrelated to MASH (e.g., hyperoxaluria, cholecystectomy, hernia repair). Written informed consent was obtained from all participants, and the study protocol received approval from the Mayo Clinic Institutional Review Board (IRB n. 21-009176) prior to initiation.

### Human Bulk and Single Cell RNA-Sequencing

Bulk human RNA-sequencing dataset GSE164760 was used to identify differentially expressed genes (DEGs) between normal livers (n=6), MASH (n=75) and MASH-HCC (n=52). Hierarchical heatmap of DEG was generated by Mayo Clinic ClustVis software. Single cell RNA (scRNA)-sequencing datasets GSE151530 and GSE125449 were downloaded from NCBI GEO and subset to samples previously categorized as HCC. The groups of samples from each dataset were processed individually following standard Seurat (v4)^12^ processing suggestions using SCTransform v2 to normalize and regress out the percentage of mitochondrial reads and cell cycle phase. These data were then integrated using Harmony (v1.2.0)^13^ and a clustering resolution of 0.4 was selected after assessment of cluster stability via clustree (0.5.1)^14^. Clusters were annotated to cell types through a combination of predictions using ScType^15^, and previously defined annotations where the highest proportion of predicted annotations per cluster were utilized to provide a cluster annotation. Selected genes visualized via feature plot are shown using a minimum expression cutoff of 1% and a maximum expression cutoff of 99%. Pearson correlation values between gene pairs were calculated by FeatureScatter in Seurat within individual clusters. All analyses were performed in R 4.1.2^16^.

### RNA Extraction and Quantitative Real-Time PCR Analysis

Total RNA was isolated from tissue using PureLink RNA Mini Kit (Thermo Fisher Scientific, 12183025). Complementary DNA (cDNA) was synthesized from 1 µg of total RNA using High-Capacity cDNA Reverse Transcription Kit (Thermo Fisher Scientific, 4368813). Quantitative real-time PCR (RT-qPCR) was conducted on a QuantStudio VI instrument (Life Technologies) using Itaq Universal SYBR green master mix (Bio-Rad, 1725124). Each sample was tested in duplicate. To determine the fold-change in gene expression compared to a control group, ΔΔCt was calculated. To determine the relative expression of corresponding genes, ΔCt was determined. A list of primers used in this study is provided in Suppl. Table 1.

### Visium Spatial Transcriptomics Analysis

Spatial transcriptomics analysis was performed with 10X Visium platform for FFPE Gene Expression Kit for Human Transcriptome (10X Genomics, 1000338), according to the manufacturer’s instructions. Sample preparation, tissue staining, imaging, library preparation and sequencing were performed as previously described^11^.

Human MASH, MASH-HCC and healthy liver samples were collected from male and female subjects (4 normal livers, 8 MASH-HCC samples) and sequenced in two different batches. Samples from each patient were in separate Visium slides. Visium slides were filtered of low-quality spots using RNA counts and a minimum feature number per spot. Samples were manually annotated as “tumor”, “non-tumor” or “healthy” based on pathological annotations. Analysis was performed in several separate ways using the pathological annotations and possible marker genes: a) merging all data and regressing sex and batch in the normalization step, followed by analysis of gene expression; b) merging all data from batch 1 only and regressing the sex and patient covariate, followed by analysis of differential gene expression; c) analysis of each slide separately and comparison of end results across conditions. Given the high impact of the confounding variables the spatial plots were re-made using a per-sample approach. Raw counts for each sample were normalized individually using the SCTransform algorithm (Seurat). The dataset dimensionality was reduced with a principal component analysis. Only between 5 - 15 principal components were used to represent each sample, i.e. gene expression across spots within the same sample was remarkably uniform, mostly for the healthy adjacent tissue. A list of 254 genes of interest to the researcher was given. The normalized relative expression of the genes (i.e. pearson residuals) in this list was plotted on a heatmap to find the most variable genes per sample. The top-most variable genes from the list given were plotted as a Spatial Feature plot to show their distribution on the tissue. Samples from batch 1 were merged and normalized using the SCTransform algorithm regressing out the patient covariate (which includes sex, age and individual variability effects). This approach was the best option found to describe the data controlling for undesired (confounding) variability. When using all data healthy “controls” were completely confounded with batch. With this normalization approach, a differential gene expression test was performed as tumor vs. non-tumor. The DEGs were tested for overrepresentation of the Hallmark gene sets from MsigDB. The gene sets were filtered to contain only genes detected in the experiment. Significance threshold for the p-value was 0.05 and multiple testing was corrected with the Benjamini and Hochberg method.

### Histopathology and Immunohistochemistry

Liver and tumor tissues were fixed in 10% formalin, embedded in paraffin (FFPE), and sectioned at 10 µm by the Mayo Clinic Pathology Research Core. Standard staining with hematoxylin and eosin (H&E) was used for histopathological assessments. Immunohistochemistry staining was performed as follow. Sections were rehydrated and incubated with 3% H_2_O_2_ to quench endogenous peroxidase activity, boiled with sodium citrate (pH 6.0) for antigen retrieval, and blocked with R.T.U. Animal-Free Blocker and Diluent (Vector Laboratories, cat. n. SP-5035-100) followed by incubation overnight at 4C with primary antibodies, including anti-glypican-3 (GPC3) (1:200, Novus Biologicals, cat. n. NBP2-44486), anti-ACSL4 (1:200, Invitrogen, cat. n. PA5-27137), and anti-ACSL5 (1:100, Invitrogen, cat. n. MA5-24595) diluted in blocking solution. The following day, sections were washed with 0.05% PBS-T and incubated with Mouse/Rabbit IgG VisUCyte HRP Polymer Antibody (R&D Systems, cat. n. VC002) for 30 minutes at room temperature. Sections were then incubated with ImmPACT DAB Peroxidase (HRP) Substrate (Vector Laboratories, cat. n. SK-4105) for 90 seconds, 2 and 3 minutes, respectively, and counterstained with Mayer’s hematoxylin. Subsequently, sections were dehydrated and mounted with Organo/Limonene Mount (ChemCruz, cat. n. sc-45087). A digital pathology scanner (MoticEasyScan Pro 6) and slide scan software (Motic DSAssistant) were used for imaging.

### Cell Culture

Human hepatocyte cell line THLE3 was purchased from ATCC (cat. n. CRL-11233), human hepatoma cell line Huh7 was kindly provided by Dr. Samar Ibrahim (Mayo Clinic Rochester), and human hepatocellular carcinoma cell line SNU475 was kindly provided by Dr. John A. Copland III (Mayo Clinic Florida). THLE3 were cultured in BEGM bullet Kit (Lonza, cat. n. CC3170), supplemented with 10% fetal bovine serum (FBS) (Gibco, cat. n. 10437-028) and 1% antibiotic-antimycotic (Gibco, cat. n. 15240-062). THLE3 were then seeded and maintained on fibronectin-collagen type I-coated plates. HCC cell lines were cultured in DMEM (Gibco, cat. n. 11995-065) supplemented with 10% FBS (Gibco, cat. n. 10437-028) and 1% antibiotic-antimycotic (Gibco, cat. n. 15240-062). Cells were maintained at 37°C in a humidified 5% CO_2_/95% air incubator. None of the cell lines used was found in the database of commonly misidentified cell lines maintained by ICLAC and NCBI Biosample. All cell lines tested negative for mycoplasma.

### Protein Isolation and Western Blot Analysis

Total proteins were extracted from THLE3, Huh7, and SNU475 cells using 1X RIPA buffer (Cell Signaling Technologies, cat. n. 9806) according to the manufacturer’s protocol. Protein concentration was measured with the Pierce BCA protein assay kit (Thermo Fisher Scientific, cat. n. 23225). Proteins were separated via SDS-polyacrylamide gel electrophoresis (SDS-PAGE) and transferred to a 0.2 µm nitrocellulose membrane (Bio-Rad, cat. n. 1620112). Membranes were then blocked with Bullet Blocking One (Nacalai USA, cat. n. 13779-01) for 5 minutes at room temperature before incubation with primary antibodies, including anti-ACSL4 (1:1000, Invitrogen, cat. n. PA5-27137) and anti-β-actin (1:1000, Abcam, cat. n. Ab8227-50), diluted in Signal Enhancer HIKARI Solution 1 (Nacalai USA, cat. n. NU00102-1) and left to shake at 4°C overnight. Following primary incubation, membranes were incubated for 1.5 hours at room temperature with the corresponding goat anti-rabbit (1:2000, Thermo Fisher Scientific, cat. n. 65-612-0) or goat anti-mouse (1:2000, Thermo Fisher Scientific, cat. n. PA186015) horseradish peroxidase (HRP)-conjugated secondary antibodies, diluted in Signal Enhancer HIKARI Solution 2 (Nacalai USA, cat. n. NU00102-2). Proteins were visualized by SuperSignal West Pico PLUS Chemiluminescent Substrate (Thermo Fisher Scientific, cat. n. 34580) and imaged with a BioRad ChemiDoc Imaging System.

### Cell Viability

Cell viability was assessed by seeding 1×10^4^ Huh7 or SNU475 cells per well in a 96-well plate. After 24 hours of incubation at 37°C/5% CO_2_, the plate was washed with PBS and 100 µL/well of fresh media was added. Cells were pre-treated with Ferrostatin-1 (1 µM, Cayman Chemical, cat. n. 17729) for 1 hour, followed by incubation with RSL-3 (5 µM, Selleck Chemicals, cat. n. S8155) for 24 hours. At the end of the study, CellTiter-Glo (Promega, cat. n. G7572) was used to assess cell viability. Luminescence was measured with a BioTek Microplate Reader.

### Animal Studies

Male, 7-9-week-old C57BL/6J mice (The Jackson Laboratory, cat. n. 000664) were housed five per cage in a 12h light-12h dark cycle. All procedures were performed according to protocols approved by the Mayo Clinic Animal Care and Use Committee (protocol n. A00004717-19-R22). Mice were fed a normal chow diet or Western Diet (WD) containing 21.1% fat, 41% Sucrose and 1.25% cholesterol by weight (Teklad diets, TD.120528). Normal water or high sugar solution containing 23.1 g/L d-fructose (SigmaAldrich, cat. n. G8270) and 18.9 g/L d-glucose (Sigma-Aldrich, cat. n. F0127) were provided. In addition, CCl_4_ (Sigma-Aldrich, cat. n. 289116) or vehicle was injected intraperitoneally weekly at the dose of 0.2 µl (0.32 µg)/g body weight, as previously described^17^. Mice were euthanized after 24 weeks by CO_2_ inhalation. Liver samples were collected and processed for further analyses.

### Murine Single Cell RNA-Sequencing

Freshly isolated livers from normal and MASH-HCC mice (n=2 mice/group) were pooled and dissociated with the tumor dissociation kit (Miltenyl Biotec, 130-096-730), according to the manufacturer’s instructions. Sample and library preparation and sequencing were performed as previously described^11^. Seurat v5 was utilized for the bioinformatic analysis of scRNA-sequencing data. Specifically, the normal liver and MASH-HCC samples were integrated and normalized. The data was then clustered, and cell markers for each cluster were identified. All cell types were manually reviewed and annotated. Subsequently, the cell counts for each cell type were compared. Finally, a dot plot was generated to illustrate both the expression levels and the percentage of cells within each cluster expressing the ACSL4 and ACSL5 genes.

## RESULTS

### ACSLs are differentially expressed in MASH and MASH-HCC

Lipid metabolic reprogramming is a hallmark of solid tumors and involves changes in genes linked to lipid metabolism. To identify lipid metabolism genes uniquely expressed in MASH-HCC as compared to non-cancerous adjacent MASH, we interrogated publicly available human bulk RNA-sequencing dataset, which included 6 normal livers, 74 MASH and 53 MASH-HCC samples. Several genes involved in lipid metabolism were identified and were differentially expressed among the three groups. However, a family of six acyl-CoA synthetase long-chain (ACSLs) enzymes emerged as a unique and differentially expressed gene family (Fig. 1A). ACSLs catalyze the conversion of long-chain fatty acids into acyl-CoA to facilitate their utilization in membrane phospholipid biosynthesis and cellular lipid metabolism. The role of ACSLs in MASH-HCC remains poorly understood. We found that ACSL4 and ACSL6, which preferentially activate long-chain polyunsaturated fatty acids (PUFA) into acyl-CoA, were greatly abundant in MASH-HCC, as compared to adjacent non-cancerous MASH and normal livers (Fig. 1A). Notably, ACSL4 has been identified as a driver of ferroptosis, a novel iron-dependent cell death mode driven by phospholipid peroxidation^18,19^. Oppositely, ACSL1 and ACSL5 were downregulated in MASH-HCC as compared to MASH and normal liver. ACSL5, which catalyzes the conversion of saturated (SFA) and monounsaturated fatty acids (MUFA), was identified as a key factor in differentiating between MASH and MASH-HCC (Fig. 1A). Based on ACSL5 preference for SFA and MUFA, it may function as a ferroptosis suppressor^20^. A pan-cancer ACSLs expression profile using ATCG data confirmed a strong upregulation of ACSL4 and downregulation of ACSL5 in HCC, as compared to normal liver (SFig. 1).

**Figure 1.**
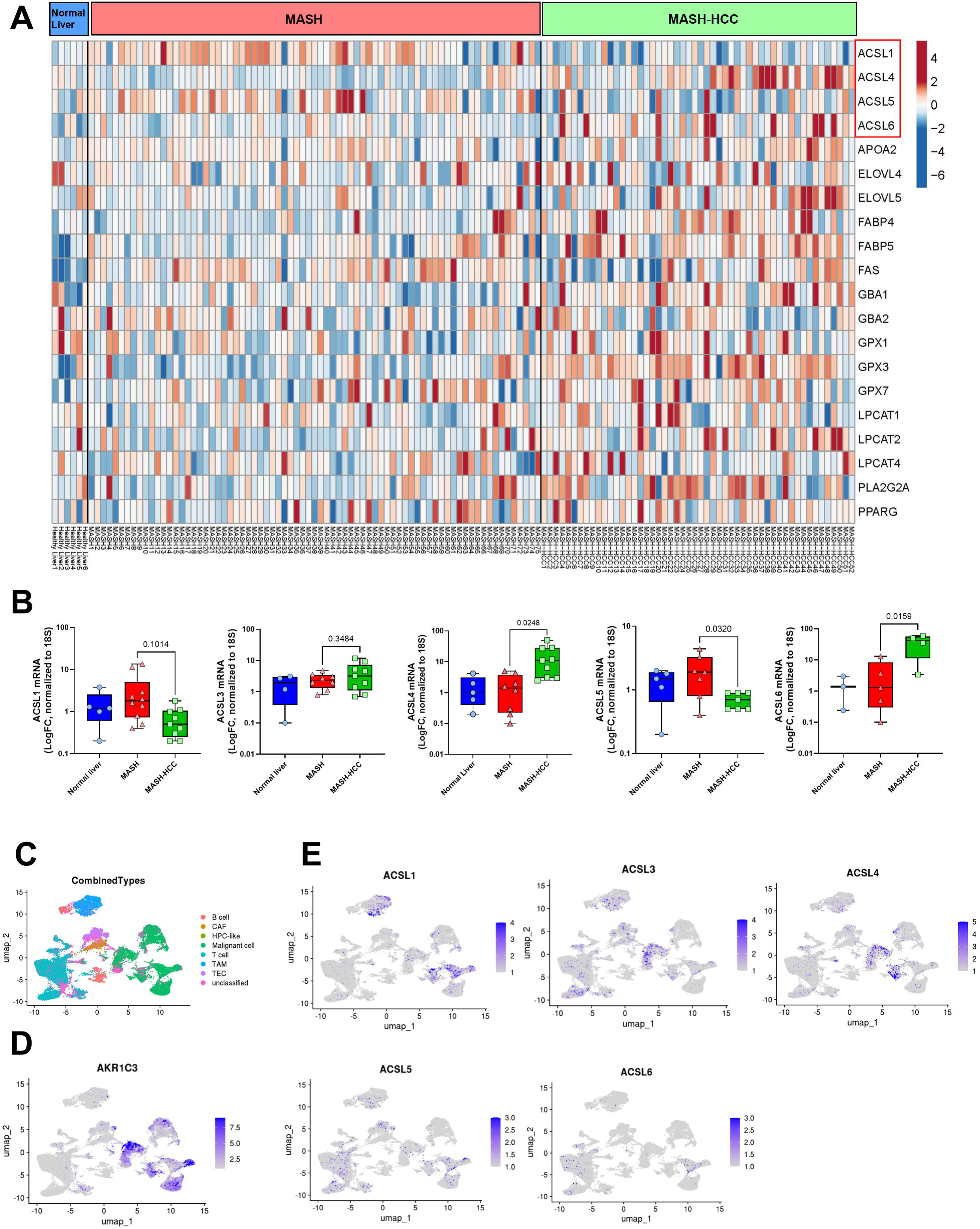
ACSLs are differentially expressed in MASH and MASH-HCC. (**A**) Heatmap of gene expression levels in normal livers, MASH, and MASH-HCC samples, analyzed from open-source bulk RNA-sequencing dataset (GSE164760). (**B**) RT-qPCR of ACSL mRNA expression in normal human livers, MASH, and MASH-HCC samples. 18S was used as a housekeeping gene. Data are presented as mean±SD. One-way ANOVAs were used for statistical analysis. (**C**) UMAP plot of liver/tumor cell clusters identified by single-cell RNA-sequencing analysis across two open-source human datasets (GSE151530, GSE125449). (**D**) UMAP plot of malignant cell marker AKR1C3. (**E**) UMAP plots of ACSL gene expression patterns.

These findings were biologically validated by RT-qPCR in a validation cohort of human normal livers (n=5 subjects), MASH (n=9 subjects) and MASH-HCC (n=9 subjects) samples. Our internal validation confirmed a strong and statistically significant upregulation of ACSL4 and ACSL5 downregulation in MASH-HCC, as compared to non-cancerous MASH (Fig. 1B). To delineate the expression of ACSLs at single-cell level, we integrated and interrogated two publically available human single-cell RNA-sequencing datasets of MASH-HCC.

The individual transcriptomes were projected on a uniform manifold approximation and projection (UMAP) plot and organized into eight distinct cell clusters (Fig. 1C). Malignant cells were defined based on expression of aldo-keto reductase family 1 member C3 (AKR1C3)^21^ (Fig. 1D). The scRNA-sequencing data analysis identified ACSL4 as highly abundant in malignant cells with strong expression peaks in selective subpopulations of malignant cells, as compared to non-cancerous cell populations (Fig. 1E). On the other hand, ACSL5 was broadly expressed in non-cancerous cell populations and almost not expressed in malignant cell populations (Fig. 1E). While ACSL6 was significantly upregulated in MASH-HCC, based on our RT-qPCR analysis (Fig. 1B), its expression was low in scRNA-sequencing data and it is mostly involved in fatty acid metabolism in brain^22^, thus for the rest of our report, we focus on the characterization of ACSL4 and ACSL5. Our findings seem to indicate that ACSLs expression is differentially regulated in MASH-HCC, where lipid metabolic reprogramming may selectively promote the expression of specific ACSLs to satisfy cancer cell metabolic needs.

### Distinct spatial and gene expression patterns of MASH-HCC

To identify region-specific lipid metabolic signatures and comprehensively map the heterogeneity of MASH-HCC, we collected 12 human tissue samples, including MASH-HCC paired with adjacent non-cancerous MASH (n=8 samples) and normal liver (n=4 samples), which were sequenced via 10X Genomics Visium platform. To assess MASH-HCC spatial diversity, tissue spots from various sections of each subject were combined and analyzed through clustering according to tissue region annotations (SFig. 2). The cluster distribution is shown in the UMAP graph (Fig. 2A). As reported in Fig. 2A, we found that while distribution of clusters corresponding to normal liver was divergent from the rest of the clusters, MASH and MASH-HCC clusters were generally intertwined, with only two clusters showing a unique and distinct distribution. Gene expression of the combined tissue regions was assessed to identify DEGs in MASH-HCC as compared to adjacent non-cancerous MASH. Our analysis revealed that key HCC markers, including glypican-3 (GPC3) and SPINK1, were upregulated in MASH-HCC clusters. Among the top upregulated markers in MASH-HCC we identified ACSL4 (Fig. 2B). A further pathway enrichment analysis based on DEGs in MASH and MASH-HCC, revealed that MASH showed enrichment of epithelial-mesenchymal transition and hypoxia pathways, while MASH-HCC exhibited enrichment of oxidative phosphorylation, mTORC1 and fatty acid metabolism pathways (Fig. 2C). Because of our interest in ACSLs, we investigated the interior spatial heterogeneity of ACSL4 and ACSL5 in the normal liver, MASH and tumor regions. We confirmed that ACSL4 is highly and predominantly expressed in the tumor regions while ACSL5 is enriched in the non-cancerous MASH regions. However, ACSL4 expression was highly heterogeneous from tumor to tumor (Fig. 2D). In the same samples, GPX4, an antioxidant enzyme that protects cells from ferroptosis, was widely expressed in all three tissue regions, but with a greater enrichment in MASH-HCC expressing high levels of ACSL4 (Fig. 2D). These findings seem to suggest that while MASH-HCC may be metabolically dependent on PUFA metabolism through ACSL4 enrichment, this may establish a ferroptosis sensitivity that could be exploited therapeutically by using GPX4 inhibitors.

**Figure 2.**
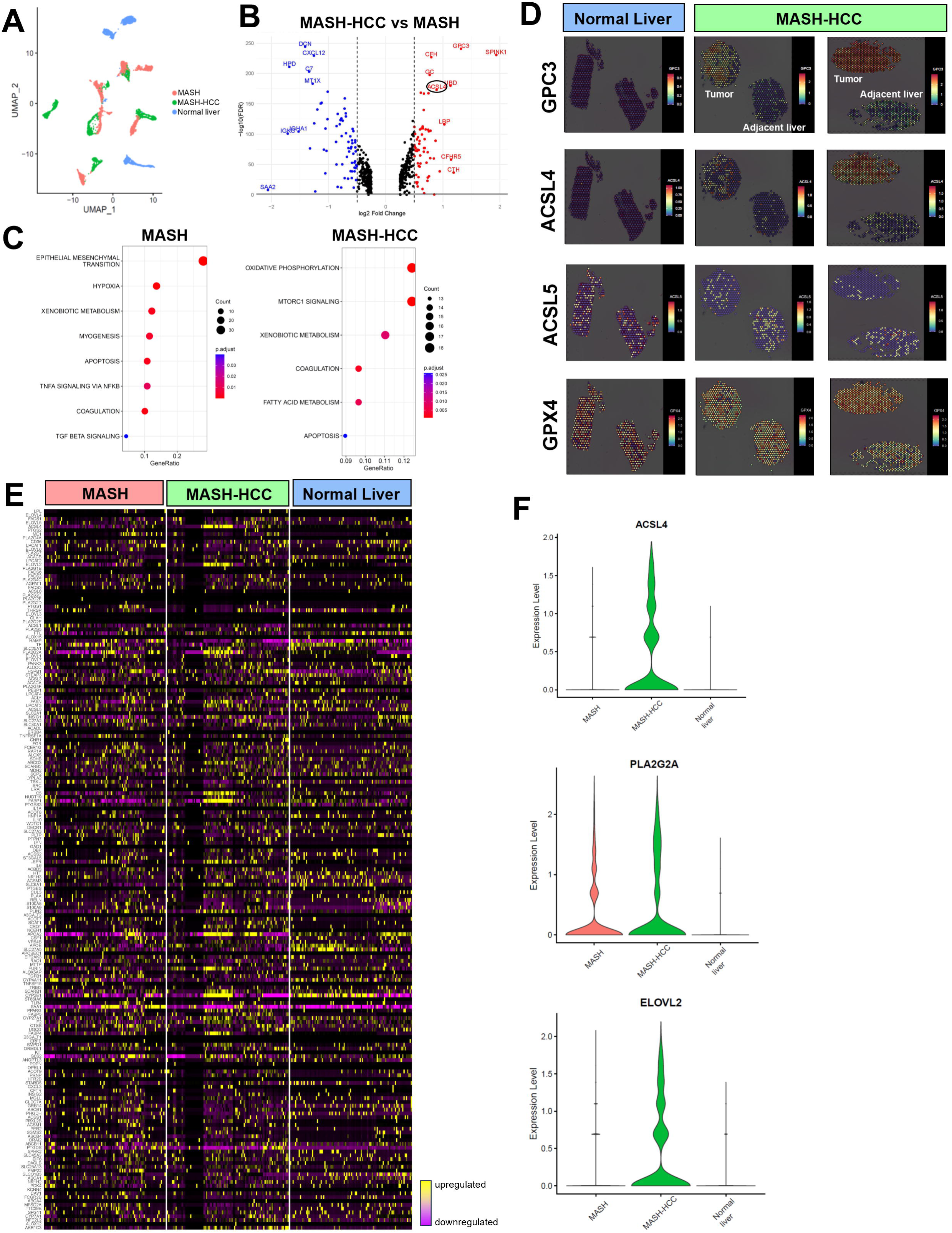
Distinct spatial and gene expression patterns of MASH-HCC. (**A**) UMAP plot of liver/tumor cell clusters identified by 10X Genomics Visium analysis. (**B**) Volcano plot of differentially expressed genes in MASH-HCC vs. MASH tissues identified by spatial transcriptomics. A log2 fold-change of 0.5 and p-value of 0.05 were used to identify statistically significant genes. (**C**) Gene sets enriched in the indicated experimental groups. (**D**) 10X Genomics spatial distribution of HCC marker GPC3, as well as ACSL4, ACSL5, and GPX4 in human MASH-HCC tissue, paired adjacent non-cancerous tissue, and normal liver tissue. (**E**) Heatmap of gene expression levels in normal, MASH, and MASH-HCC tissues. (**F**) Violin plots of lipid metabolism genes identified as upregulated in MASH-HCC compared to MASH tissues.

To further expand our characterization of MASH-HCC diversity, we profiled the expression of genes involved in lipid metabolism in the three tissue regions, normal liver, adjacent non-cancerous MASH and MASH-HCC. A hierarchical clustering and associated violin plots reveal distinct clusters of genes with differential expression across tissue regions. Specifically, genes such as ELOVL2, PLA2G2A, FABP1, APOA2, LPCAT3, FABP4 and FABP5 were enriched in MASH-HCC, as compared to non-cancerous MASH and normal liver regions (Fig. 2E-F). Among these genes, FABP5 was recently proposed as a biomarker of ferroptosis in hypoxic brain^23^. These findings indicate that spatial intratumor heterogeneities are common. The various tissue clusters exhibit diverse pathway activities and lipid metabolic gene signatures. Studying the differential lipid metabolism profiles may identify metabolic vulnerabilities to target for novel MASH-HCC treatments.

### ACSL4 is a biomarker of MASH-HCC and is associated with ferroptosis sensitivity in HCC cells

To further validate our spatial transcriptomics findings, we performed targeted immunohistochemistry characterization in larger tissue specimens, consisting of adjacent non-cancerous MASH and MASH-HCC regions on the same tissue section. We collected 3 new human MASH-HCC tissue samples which were paired with their respective hematoxylin-eosin labeling and stained for HCC marker GPC3 to differentiate the tumor region from the non-cancerous adjacent area (Fig. 3A). Immunohistochemistry on the validation specimens identified ACSL4 highly expressed in GPC3-positive tumor regions, whereas ACSL5 was mostly enriched in non-cancerous tissue (Fig. 3A). Despite this tissue region selectivity of ACSL4 and ACSL5, both markers showed sample-to-sample heterogeneity, supporting our spatial transcriptomics profiling.

**Figure 3.**
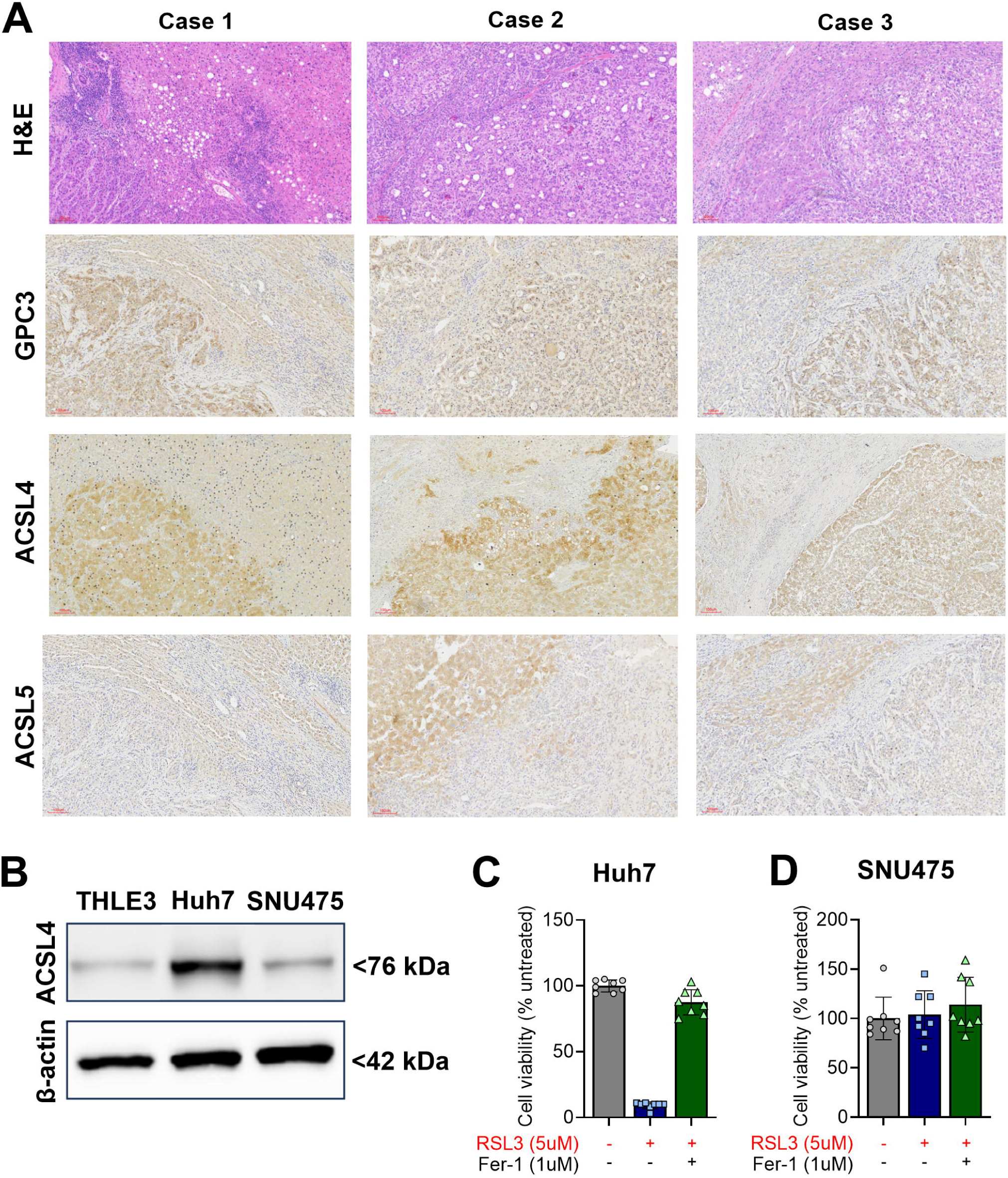
ACSL4 is a biomarker of MASH-HCC and is associated with ferroptosis sensitivity in HCC cells. (**A**) Histopathological and immunohistochemistry staining of three human MASH-HCC subjects, including H&E staining (top), GPC3, ACSL4 and ACSL5. (**B**) Western blot of ACSL4 protein expression in normal human hepatocytes (THLE3) and HCC cell lines (Huh7, SNU475). β-actin was used as a loading control. (**C-D**) CellTiter-Glo cell viability assessment in Huh7 and SNU475 HCC cell exposed to ferroptosis inducer RSL3 (5µM) for 24 hours and rescued by lipid peroxidation inhibitor ferrostatin-1 (1µM). One-way ANOVA was used for statistical analysis.

We then sought to investigate the expression of ACSL4 in normal human hepatocytes (THLE3) and two HCC cell lines (Huh7 and SNU475) to test *in vitro* whether ACSL4 protein expression is associated with sensitivity to ferroptosis. First, ACSL4 was highly expressed in Huh7 but mildly expressed in normal hepatocytes and SNU475, confirming both the enrichment of ACSL4 and its expression heterogeneity in HCC cells (Fig. 3B). Second, both Huh7 and SNU475 were treated with ferroptosis inducer RSL3, an inhibitor of endogenous antioxidant GPX4, which protects cells from ferroptosis. RSL3-induced ferroptosis was abrogated by co-treatment with ferrostatin-1, a synthetic scavenger that protects cells from RSL3-induced lipid peroxidation and ferroptosis. Cell viability assessed by CellTiter-Glo shows a dramatic 90% loss of cell viability of Huh7 cells treated with RSL3 for 24 hours, as compared to DMSO-treated control cells (p<0.0001). RSL3-induced Huh7 cell viability loss was rescued by addition of ferroptosis inhibitor ferrostatin-1 (Fig. 3C). On the contrary, SNU475 cells treated with ferroptosis-inducer RSL3 exhibited complete ferroptosis resistance, which might be a consequence of the significantly lower expression of ACSL4 (Fig. 3D). Our findings suggest that ACSL4 may serve as an *in vivo* biomarker of ferroptosis in MASH-HCC and may predict ferroptosis sensitivity, thus providing an opportunity to improve the efficacy of current standard-of-care HCC therapy.

### Transcriptional profiling of ACSL4 and ACSL5 in an experimental murine model of MASH-HCC

Our findings thus far focus on cancer cells and hepatocytes. However, stromal and immune cells play a critical role in MASH and MASH-HCC progression. In addition, the role of ACSLs in these cells is currently poorly understood. To characterize ACSL4 and ACSL5 in other cell populations, we performed a single-cell RNA-sequencing analysis in an experimental model of diet-induced MASH-HCC. Mice were fed a western diet (WD) and received a weekly injection of hepatotoxin carbon tetrachloride (CCl_4_) for 24 weeks to develop MASH-HCC, as compared to control mice which were fed a chow diet (Fig. 4A). Gross liver images confirmed the development of nodules (yellow circles) in the MASH-HCC mouse group as compared to the control group, where livers appear healthy (Fig. 4B). Further histopathological examination of harvested livers showed MASH development in mice fed a western diet, with presence of liver steatosis and inflammation, as compared to normal liver tissue in control mice (Fig. 4C). The nodules spotted in our gross images were diagnosed as dysplastic nodules with steatohepatitic features. Cell clusters from both experimental mouse groups were overlapped and reported in a UMAP plot (Fig. 4D). Additional uniform manifold approximation and projection dimensionality reduction analysis generated combining data from both experimental groups, identified 18 cell clusters that were appropriately annotated based on cell-specific enriched genes (SFig. 3). The clusters corresponded to the main liver cell populations, including epithelial cells (hepatocytes, cholangiocytes), innate and adaptive immune cells, including those associated with the tumor immune microenvironment and stromal cells (hepatic stellate cells, liver sinusoidal cells). We also identified a model-specific cluster of dysplastic cells in the MASH-HCC livers (Fig. 4E). Normal and MASH-HCC livers showed differential abundance of each cell populations. MASH-HCC livers exhibited larger amount of innate and adaptive immune cells, including monocyte-derived macrophages, B cells, CD8+ T cells, dendritic cells (DC), T regulatory cells (Treg), natural killer cells (NK) and natural killer T cells (NKT), as compared to normal livers (Fig. 4F). As expected, dysplastic malignant cells were identified only in the MASH-HCC livers while quiescent hepatic stellate cells (qHSC) were identified only in the normal livers (Fig. 4F). We then performed a bioinformatics analysis to identify the differential expression of ACSL4 and ACSL5 in all the cell populations identified, which were grouped by cell type in epithelial, immune and stromal cells (Fig. 4G). Along with our previous findings, malignant dysplastic cells showed higher expression of ACSL4 than ACSL5, which was more abundantly expressed in non-dysplastic epithelial cells, such as cholangiocytes. Unexpectedly, we found a 2-fold upregulation of ACSL4 in NKT and NK cells from the MASH-HCC livers, as compared to those from normal livers (Fig. 4G). With the exception of dendritic cells, ACSL4 expression was enriched in immune cells in the MASH-HCC livers, as compared to normal livers. Our analysis also found ACSL5 particularly enriched in endothelial cells, independently of the experimental group. Our findings seem to point to a potential key role of ACSL4 in shaping MASH-HCC immune microenvironment and ACSL5 in regulating stromal and non-cancerous cell biology. These findings warrant further investigation to uncover the specific roles of ACSLs in tumor immunity and whether some of these functions may be targeted for better MASH-HCC treatments.

**Figure 4.**
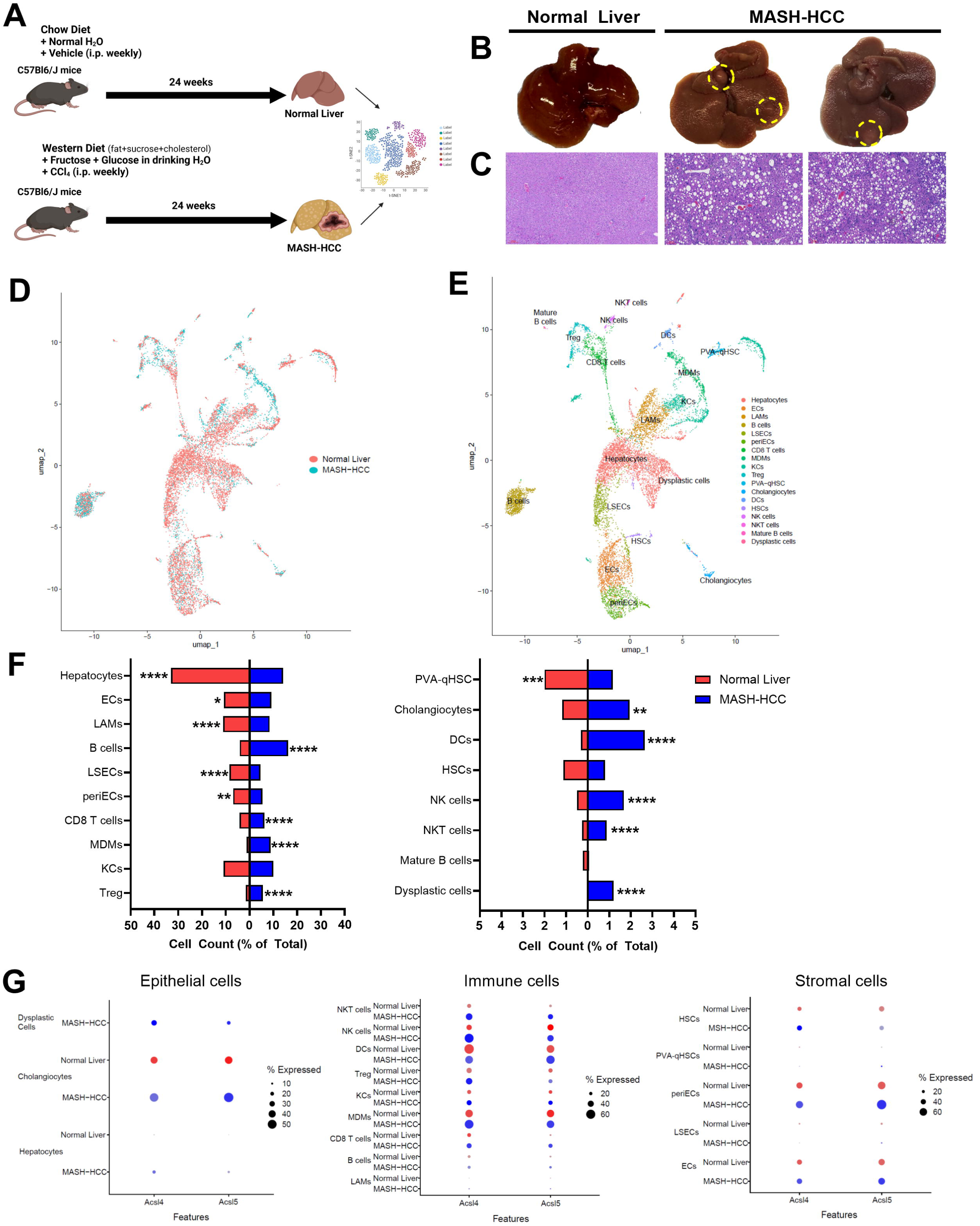
Transcriptional profiling of ACSL4 and ACSL5 in an experimental murine model of MASH-HCC. (**A**) Graphical representation of diet-induced MASH-HCC preclinical mouse model. Mice were fed a Western diet plus sugar water and injected weekly with CCl_4_ intraperitoneally for 24 weeks. (**B**) Gross images of livers highlighting the formation of dysplastic nodules (yellow circles) in the indicated experimental groups. (**C**) Representative H&E staining of liver tissues for the indicated experimental groups. (**D**) UMAP plot of liver/tumor cell transcriptomes colored according to group (n= 2 mice/group). (**E**) UMAP plot of liver/tumor cell clusters colored according to their identified cell type. (**F**) Relative cell count normalized on total count of each cell cluster in the indicated experimental groups. A two-sample proportion test was used for statistical analysis. \**p*<0.05, \*\**p*<0.01, \*\*\**p*<0.001, \*\*\*\**p*<0.0001. (**G**) Expression levels and percentage of cells within each cluster expressing the ACSL4 and ACSL5 genes in the indicated experimental groups.

## DISCUSSION

Metabolic reprogramming is a hallmark of solid tumors and is essential for tumor growth and survival, especially in metabolically stressed conditions^24^. Among metabolic alterations in cancer, dysregulation of lipid metabolism is the most prominent^6^. In a lipid-rich environment such as that of MASH, exogenous and endogenous fatty acids represent a key source for mitochondrial and peroxisomal fatty acid oxidation (FAO), phospholipid and cholesteryl esters synthesis for membrane biogenesis, especially under rapid proliferative states^7,25^. Before they can be utilized in the aforementioned pathways, intracellular short-chain, medium-chain and long-chain fatty acids (LCFA) are converted into fatty acyl-CoA by acyl coenzyme A synthetase in an ATP-dependent reaction. Long-chain acyl-CoA synthetases (ACSLs) are key members of the acyl coenzyme A synthetases and prefer fatty acids with 12 to 20 carbon atoms^26^. Notably, each member of the ACSL family catalyzes the activation of different LCFAs which after being converted to LCFA-CoA undergo self-modification, elongation and desaturation, thus fueling various catabolic and energy-producing metabolic pathways.

ACSL1 is predominantly expressed in the liver and abundantly localized on lipid droplets, microsomes and mitochondria^27,28^. It is involved in fatty acid intake by activating membrane fatty acid transporters, FAO and triglyceride synthesis. ACSL1 prefers oleate and linoleate acids for activation^29^. ACSL3 is highly abundant in brain and is involved in fetal development, lipid droplet formation and *de novo* lipogenesis^30^. ACSL3 preferentially incorporates myristate, laurate, arachidonate and eicosapentaenoate into phosphatidylcholine, the major very low-density lipoprotein (VLDL) phospholipid^31^. ACSL4 prefers PUFAs, such as arachidonate and eicosapentaenoate, as substrates and it is involved in the synthesis of lipid mediators such as prostaglandin E2^32^. ACSL5 exhibit a preference for a wide range of saturated fatty acids, such as palmitate, palmitoleate and C16-18 unsaturated fatty acids, such as oleate. ACSL6 plays a crucial role in phospholipid and triglyceride synthesis in brain and prefers C16-C20 saturated and polyunsaturated fatty acids^33^.

Our report aimed at identifying lipid gene signatures between MASH-HCC and non-cancerous MASH, reveals a differential expression of ACSL4 and ACSL5 at the transcriptional, spatial and single-cell level. The landscape of lipid metabolic alterations driving malignant transformation and progression of MASH-HCC remains poorly understood and a key research priority for this tumor. Our integrated analysis shed lights on the expression pattern of ACSLs in MASH-HCC. This is relevant because ACSLs not only contribute to tumor proliferation, metastatic activity and drug resistance but may also constitute a metabolic vulnerability by conferring cancer cells sensitivity to cell death. An example of cell death tightly linked to lipid metabolism is ferroptosis.

Ferroptosis is a novel mode of iron-dependent and caspase-independent cell death driven by phospholipid peroxidation^8,34^. Ferroptosis occurs as a consequence of unabated redox-active iron accumulation in cells and subcellular organelles, failure to detoxify highly reactive lipid peroxides and membrane phospholipid peroxidation causing membrane rupture, the point of no return in ferroptosis^10^. Intriguingly, if the amount of PUFA in membrane phospholipids determines membrane fluidity, it also increases the intrinsic susceptibility to ferroptosis. Therefore, the balance between ferroptosis-blocking saturated or monounsaturated fatty acid-phospholipids vs. ferroptosis-inducing PUFA-phospholipids dictates ferroptosis sensitivity.

In cancer, ferroptosis can either inhibit or advance tumor growth, depending on the cell, tumor type and tumor immune microenvironment. Ferroptotic cancer cells may release signals and lipid mediators that can exert either immunogenic or immunosuppressive functions^35^. Ferroptosis has been shown to promote MASH progression through impairment of GPX4, a major anti-ferroptosis defense mechanism^36^. Ferroptosis has been explored also as a new treatment for HCC. However, it was found that HCC rewires its metabolism to escape ferroptosis^37^. Therefore, identifying which lipid metabolic markers are upregulated and co-expressed in MASH-HCC may uncover novel biomarkers of *in vivo* ferroptosis and may provide insights on mechanisms that regulate ferroptosis sensitivity in immunocompetent systems.

ACSL4 is upregulated in several solid tumors and ACSL4-driven PUFA metabolism has been associated with ferroptosis sensitivity^18,38,39^. In pre-clinical murine models of MASH-HCC, ACSL4 upregulation has been linked to the disease progression through enhanced lipid accumulation and inflammation^40^. ACSL4 was found to execute its role as determinant of ferroptosis sensitivity by remodeling membrane phospholipid composition with ferroptosis-inducing PUFA phospholipids^18^. Our integrative report found that overall ACSL4 is enriched in MASH-HCC as compared to non-cancerous MASH and healthy livers. However, intratumor ACSL4 expression is highly heterogeneous, suggesting that while some MASH-HCC may rely on PUFA metabolism for their metabolic demand, thus acquiring a metabolic vulnerability, others rely on different metabolic pathways, thus may acquire ferroptosis resistance. Given the complexity of lipid metabolism, ACSL4 does not work alone. We found that genes involved in lipid metabolism are co-expressed and co-localize with ACSL4 in MASH-HCC. These additional genes included ELOVL2, PLA2G2A, FABP1, APOA2, FABP4, LPCAT3, AKR1C3 and G0S2 which may provide a more reliable gene signature of ferroptosis sensitivity in high ACSL4-expressing MASH-HCCs. In support of these findings, our *in vitro* studies revealed that in human HCC cells, ACSL4 protein expression correlated with ferroptosis sensitivity. HCC cells expressing low amount of ACSL4 were resistant to RSL3-induced ferroptosis, which was rescued by ferroptosis inhibitor ferrostatin-1.

Particularly notable is the finding that ACSL5 seems to function in opposition to ACSL4. We found that ACSL5 is more abundant in non-cancerous MASH and healthy liver tissues than MASH-HCC. While the role of ACSL5 has not been well-established in ferroptosis, the preference of ACSL5 for saturated and monounsaturated fatty acids as substrates, classify this ACSL as a potential ferroptosis resistance marker. The function of ACSL5 in MASH-HCC is currently poorly understood. Its enrichment in non-cancerous MASH tissue and normal liver seem to indicate that ACSL5 is crucial for maintaining lipid homeostasis. We also showed that ACSL4 and ACSL5 have a preferential expression in different cell populations of MASH-HCC based on our single-cell RNA-sequencing analysis of murine normal liver and MASH-HCC. In experimental models of peritonitis, ACSL4-deficiency in peritoneal macrophages reduced inflammation and biosynthesis of lipid mediators such as leukotriene B4 and PGE2, which may have tumor-promoting or suppressive functions^41^.

Evidence points to a key role of tumor ACSLs and their LCFA substrates in modulation of tumor immune responses. However, the functions of ACSLs in immune cells remain poorly understood. While we did not perform a functional analysis of ACSLs in immune cells, we report single-cell transcriptomics data on the expression of ACSL4 and ACSL5 across multiple cell populations of murine normal and MASH-HCC livers. Overall, we confirmed that ACSL4 is preferentially enriched in dysplastic malignant cells as compared to normal epithelial cells, such as hepatocytes and cholangiocytes. Intriguingly, we observed a marked enrichment of ACSL4 in almost all immune cells of MASH-HCC, particularly in NK and NKT cells, suggesting that ACSL4 may play a crucial role in shaping the MASH-HCC immune microenvironment.

How ACSL4 functions in immune cells such as NK and NKT cells and how ACSL4 modulation reprograms antitumor immunity and regulate ferroptosis sensitivity remain open questions that will be investigated in a follow up study.

In summary, we have identified and characterized ACSLs expression patterns in human normal liver, MASH and MASH-HCC using an integrated strategy that included bulk RNA-sequencing, scRNA-sequencing and spatial transcriptomics analyses. These studies not only identified ACSL4 and ACSL5 with opposite functions and expression profiles but may have uncovered a gene signature of genes involved in lipid metabolism that may be predictive of ferroptosis sensitivity *in vivo*. We validated key findings *in vitro* and by immunohistochemistry staining which suggest that ACSL4 appears to be a marker of ferroptosis sensitivity in MASH-HCC. Moreover, we profiled ACSL4 and ACSL5 expression in a multitude of cells in an experimental murine model of MASH-HCC, which revealed a potential crucial role of ACSL4 in shaping the MASH-HCC tumor immune microenvironment. Our findings encourage further investigation and mechanistic studies to better understand the role of ACSL4 and ACSL5 in MASH-HCC. These insights may also provide novel therapeutic opportunities for example by combining current inhibitors for PCSK9 and next generation GPX4 inhibitors for the treatment of HCC.

## STUDY LIMITATIONS

Our report remains mainly descriptive with only minimal mechanistic insights because we focus on a broad characterization of ACSLs in MASH-HCC. A follow-up study that will focus on how are ACSL4 and ACSL5 regulated in MASH-HCC and what is their role in tumor immunity is in our pipeline. Further spatial lipid profiling in MASH-HCC is also relevant for this project and will be included in our future studies.

## ACKNOWLEDGEMENTS

Assistance with the study: We would like to thank Dr. Lisa Boardman, Donna Felmlee-Devine and Jennifer S. Horsch, of the Mayo Clinic Center for Cell Signaling in Gastroenterology for the help with IRB request, human specimens’ procurement, infrastructure and resources needed for this study. We would like to thank Dr. John A. Copland III in the Mayo Clinic Department of Cancer Biology for kindly providing HCC cell lines. We would like to thank Dr. Kengunte Nagaraj Nagaswaroop and Dr. Yue Yu from the Mayo Clinic Department of Quantitative Health Sciences for the help with RNA-sequencing analysis. We would like to thank Vernadette A. Simon and Fariborz Rakhshan Rohakhtar of the Mayo Clinic Genomics Core for the help with library preparation and scRNA-sequencing. We would like to thank RaVuth Keo of the Mayo Clinic Pathology Research Core for the assistance with tissue sections and processing for spatial transcriptomics, Colleen Forster of the University of Minnesota Histology and Research Laboratory for guidance with tissue deparaffinization and staining, Grant Barthel of the University of Minnesota Imaging Center for performing Visium gene expression slide imaging, Fernanda Rodriguez and John Garbe of the University of Minnesota Genomics Center for spatial library preparation and transcriptome sequencing. We would like to thank Dr. Valentina Nardi of Mayo Clinic Department of Cardiology for letting us use the MoticScanner for tissue slide imaging.

## AUTHOR CONTRIBUTIONS

Conceived the idea: DP, RPG, NT; Designed the experiments: PC, DP; Performed the experiments: PC; Analyzed the data: PC, DP; Performed the bioinformatics analyses: AW, NCM; Contributed human specimens: RPG, NT, TT; Contributed reagents/materials/analysis tools: PC, DP; Wrote/revised the manuscript: PC, DP; Reviewed/edited the manuscript: All authors discussed and commented on the manuscript.

## ABBREVIATIONS

MASLD: Metabolic-dysfunction associated steatotic liver disease
MASL: Metabolic-dysfunction associated steatotic liver
MASH: Metabolic-dysfunction associated steatohepatitis
HCC: Hepatocellular carcinoma
MASH-HCC: MASH-driven hepatocellular carcinoma
PUFA-PL: Polyunsaturated fatty acid-containing membrane phospholipids
GPX4: Glutathione peroxidase 4
FSP1: Ferroptosis suppressor protein 1
ACSLs: Acyl-CoA synthetase long chain
NK: natural killer cells
NKT: natural killer T cells
IRB: Institutional Review Board
scRNA: single-cell RNA
DEGs: differentially expressed genes
cDNA: Complementary DNA
RT-qPCR: Real-time quantitative polymerase chain reaction
FFPE: Formalin-fixed paraffin-embedded
H&E: Hematoxylin and eosin
GPC3: Glypican-3
FBS: fetal bovine serum
SDS-PAGE: SDS-polyacrylamide gel electrophoresis
HRP: horseradish peroxidase
CO_2_: Carbon dioxide
WD: Western diet
CCl_4_: Carbon tetrachloride
PUFA: polyunsaturated fatty acids
SFA: Saturated fatty acids
MUFA: Monounsaturated fatty acids
UMAP: Uniform manifold approximation and projection
AKR1C3: Aldo-keto reductase family 1 member C3
GC: group-specific component
SPINK1: serine protease inhibitor Kazal type 1
ELOVL2: elongation of very long chain fatty acids-like 2
PLA2G2A: phospholipase A2 group IIA
FASN: fatty acid synthase
FABP1: fatty acid binding protein 1
APOA2: apolipoprotein A-II
LPCAT3: lysophosphatidylcholine acyltransferase 3
FABP4: fatty acid binding protein 4
RSL3: RAS-selective lethal 3
DC: Dendritic cells
Treg: T regulatory cells
qHSC: quiescent hepatic stellate cells
FAO: fatty acid oxidation
LCFA: long-chain fatty acids
VLDL: very low-density lipoprotein
G0S2: G0/G1 switch gene 2

