## Supplemental Material for "Role of Long Chain Acyl-CoA Synthetases in MASH-driven Hepatocellular Carcinoma and Ferroptosis"

**Number of Text Pages:** 23

**Number of Tables:** 1 Supplemental

**Number of Figures:** 4 main, 3 supplemental

**Running Head:** Lipid metabolism landscape and ferroptosis in MASH-HCC

**Financial Support:** This study was supported by the Mayo Clinic Center for Biomedical Discovery Pilot Award and NCI-funded Mayo Clinic SPORE in Hepatobiliary Cancer (P50 CA210964) Career Enhancement Program to DP.

**Conflict of Interest:** None to disclose

**Keywords:** Lipid Metabolism, Cell Death, Multi-omics analysis, Spatial transcriptomics, ACSL4, ACSL5

### SUPPLEMENTAL FIGURE LEGENDS

**Supplemental Figure 1. Pan-cancer expression of ACSLs.** Bioinformatic pan-cancer analysis of ACSL expression in human primary tumors and normal tissues determined by TCGA data sets. BLCA, bladder urothelial carcinoma; BRCA, breast invasive carcinoma; CESC, cervical squamous cell carcinoma; CHOL, cholangiocarcinoma; COAD, colon adenocarcinoma; ESCA, esophageal carcinoma; GBM, glioblastoma multiforme; HNSC, head and neck squamous cell carcinoma; KICH, kidney chromophobe; KIRC, kidney renal clear cell carcinoma; KIRP, kidney renal papillary cell carcinoma; LIHC, liver hepatocellular carcinoma; LUAD, lung adenocarcinoma; LUSC, lung squamous cell carcinoma; PAAD, pancreatic adenocarcinoma; PRAD, prostate adenocarcinoma; PCPG, pheochromocytoma and paraganglioma; READ, rectal adenocarcinoma; SARC, sarcoma; SKCM, skin cutaneous melanoma; THCA, thyroid carcinoma; THYM, thymoma; STAD, stomach adenocarcinoma; UCEC, uterine corpus endometrial carcinoma.

**Supplemental Figure 2. Spatial transcriptomics pathology markers.** Violin plot of top differentially expressed genes, based on adjusted p-value, for each indicated experimental group (Normal, MASH, MASH-HCC).

**Supplemental Figure 3. Mouse scRNA-sequencing annotation.** Heatmap of manual annotation of cell clusters identified by scRNA-sequencing. ECs, endothelial cells; LAMs, lipid-associated macrophages; LSECs, liver sinusoidal endothelial cells; periECs, periportal endothelial cells; MDMs, monocyte-derived macrophages; KCs, Kupffer cells; Treg, regulatory T cells; PVA-qHSC, quiescent hepatic stellate cells; DCs, dendritic cells; HSCs, hepatic stellate cells; NK cells, natural killer cells; NKT cells, natural killer T cells.

### SUPPLEMENTAL TABLE LEGEND

**Supplemental Table 1.** Sequences of primers used for RT-qPCR.

**Supplemental Figure 1. Classon et al.**

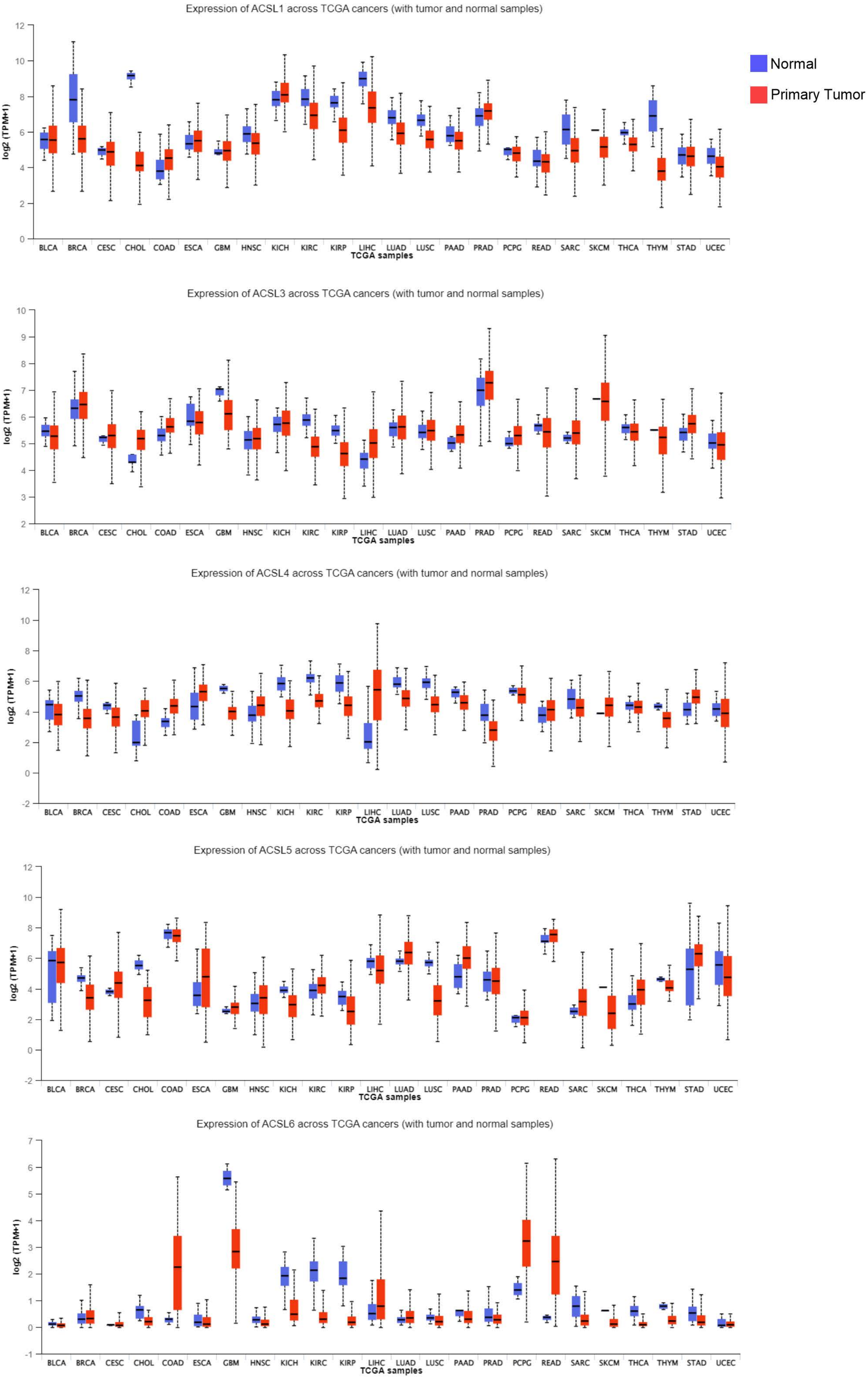

**Supplemental Figure 2. Classon et al.**

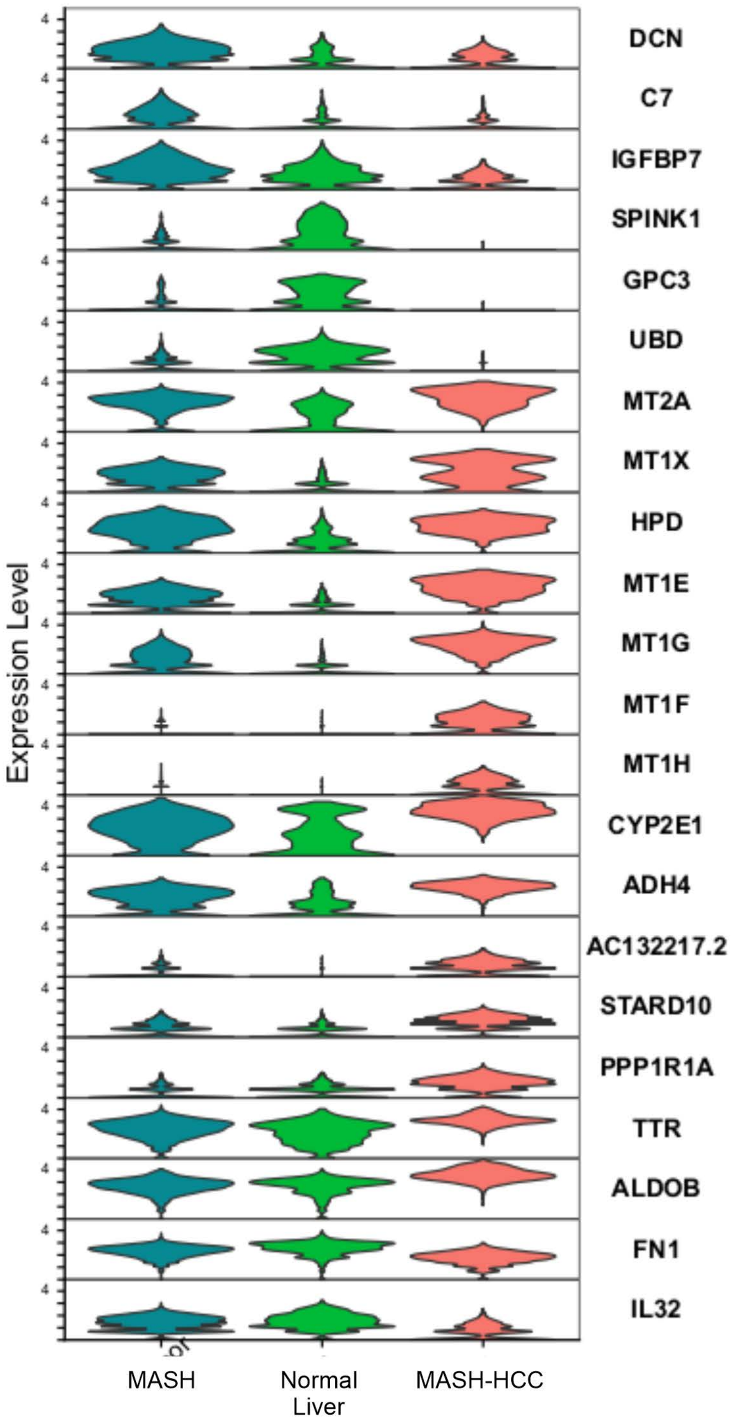

**Supplemental Figure 3. Classon et al.**

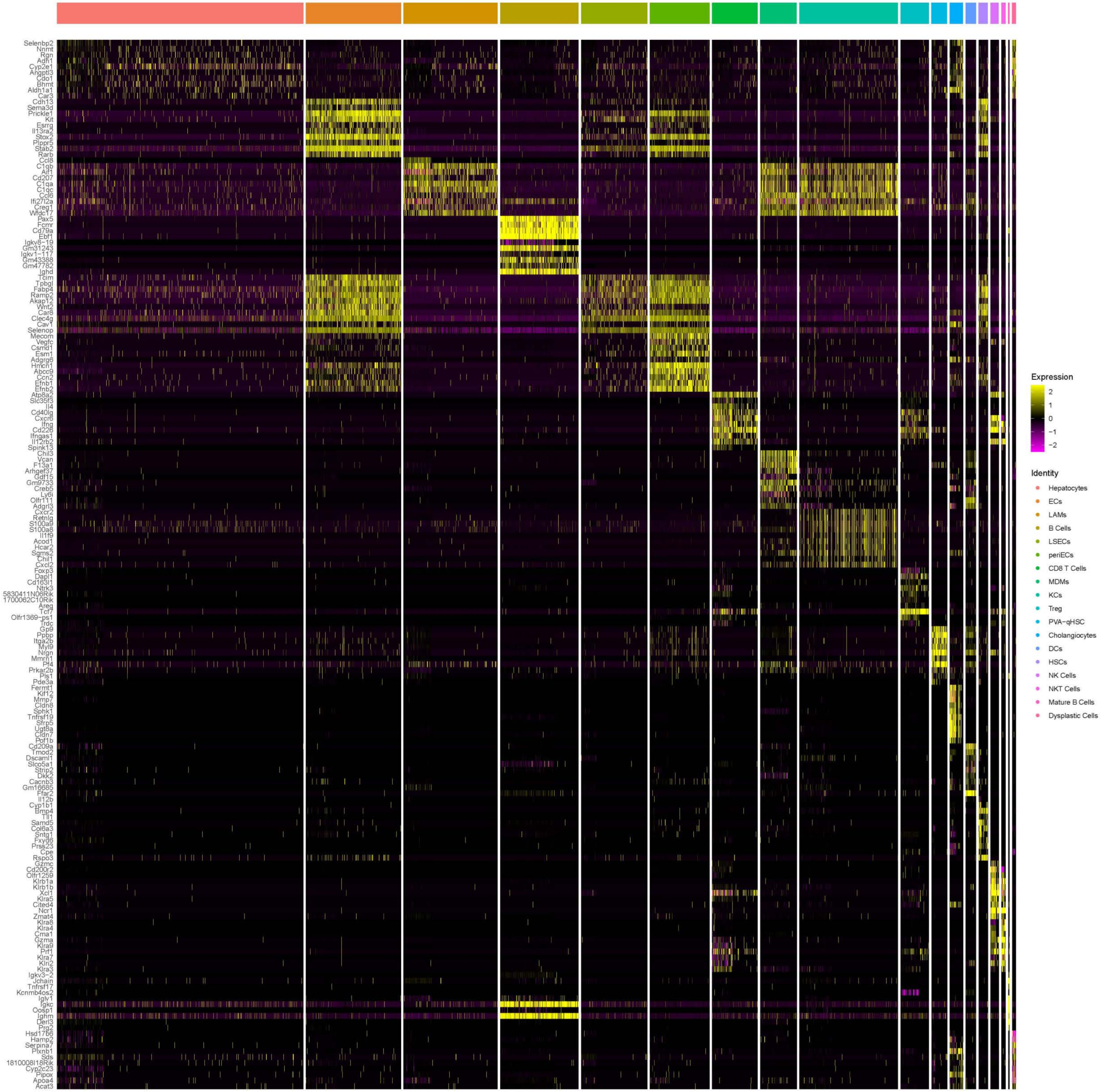

| **Target** | **Sequences** |
|---|---|
| hACSL1 | FW 5' – CCA TGA GCT GTT CCG CTA TTT – 3'<br>RV 5' – CCG AAG CCC ATA AGC GTG TT – 3' |
| hACSL3 | FW 5' – GCC GAG TGG ATG ATA GCT GC – 3'<br>RV 5' – ATG GCT GGA CCT CCT AGA GTG – 3' |
| hACSL4 | FW 5' – CAT CCC TGG AGC AGA TAC TCT – 3'<br>RV 5' – TCA CTT AGG ATT TCC CTG GTC – 3' |
| hACSL5 | FW 5' – CTC AAC CCG TCT TAC CTC TTC T – 3'<br>RV 5' – GCA GCA ACT TGT TAG GTC ATT G – 3' |
| hACSL6 | FW 5' – CGC TAC ATC AAT ACA GCG G – 3'<br>RV 5' – GCA TGG ACT TAA TGA CCA CCC – 3' |
| h18s | FW 5' – CTA CCA CAT CCA AGG AAG CA– 3'<br>RV 5' – TTT TTC GTC ACT ACC TCC CCG– 3' |
